# Effects of beta- and gamma-band rhythmic stimulation on motor inhibition

**DOI:** 10.1101/2020.12.11.422006

**Authors:** Inge Leunissen, Manon Van Steenkiste, Kirstin-Friederike Heise, Thiago Santos Monteiro, Kyle Dunovan, Dante Mantini, James P. Coxon, Stephan P. Swinnen

**Author notes:** James P. Coxon and Stephan P. Swinnen should be considered joint senior author. **Corresponding author:** Inge Leunissen, Oxfordlaan 55, 6229 EV, Maastricht, The Netherlands.

## Abstract

Voluntary movements are accompanied by an increase in gamma-band oscillatory activity (60-100Hz) and a strong desynchronization of beta-band activity (13-30Hz) in the motor system at both the cortical and subcortical level. Conversely, successful motor inhibition is associated with increased beta power in a fronto-basal-ganglia network. Intriguingly, gamma activity also increases in response to a stop-signal. In this study, we used transcranial alternating current stimulation to drive beta and gamma oscillations to investigate whether these frequencies are causally related to motor inhibition. We found that 20Hz stimulation targeted at the pre-supplementary motor area enhanced inhibition and increased beta oscillatory activity around the time of the stop-signal in trials directly following stimulation. In contrast, 70Hz stimulation seemed to slow down the braking process, and predominantly affected go task performance. These results demonstrate that the effects of tACS are state-dependent and that especially fronto-central beta activity is a functional marker for successful motor inhibition.

## Introduction

Inhibitory control, such as the ability to suppress an already initiated movement, is essential in everyday life. Successful motor inhibition activates a distributed network of cortical and subcortical areas in which the pre-supplementary motor area (preSMA), the right inferior frontal cortex (rIFC) and the subthalamic nucleus (STN) have been identified as key nodes (Aron et al., 2016; Jahanshahi et al., 2015). However, the exact nature of the neural dynamics within this fronto-basal-ganglia network are not entirely clear.

Long-distance neural communication is thought to arise from groups of neurons engaging in rhythmic synchronization (Fries, 2005). In the human motor system two main natural rhythms have been identified. On both a cortical and subcortical level gamma-band oscillatory activity (60-100Hz) increases during voluntary movement (Crone et al., 1998; Litvak et al., 2012), suggesting it has a prokinetic role. In contrast, oscillatory activity in the beta-band (13-30Hz) is prominent during tonic contractions, and decreases prior to and during movement (Engel & Fries, 2010; Schmidt et al., 2019). Excessive beta oscillations, as in Parkinson’s disease, are associated with slowing of movement and rigidity (Kuhn et al., 2004; Little & Brown, 2014). This has led to the idea that beta activity might promote the inhibition of movement.

Indeed, electrophysiological recordings have revealed increased beta oscillations in preSMA, rIFC and STN during successful motor inhibition (Alegre et al., 2013; Castiglione et al., 2019; Huster et al., 2017; Kuhn et al., 2004; Ray et al., 2012; Swann et al., 2009; Swann et al., 2012; Wagner et al., 2018; Wessel et al., 2013; Wessel et al., 2016). Crucially, this activity was seen after the presentation of a stop-signal, but before the completion of the stop process (as indexed by the stop-signal reaction time; SSRT). Yet, others reported that beta oscillations primarily increase after the SSRT (Fischer et al., 2017; Jha et al., 2015), or without differentiation between successful and unsuccessful stops (Fonken et al., 2016). These authors suggest that fronto-subthalamic beta activity is not *necessary* for stopping, but rather reflects post-processing of the stop-signal trial, and is perhaps responsible for the slowing that is typically observed on trials that follow a stop signal (Bissett & Logan, 2012). Thus controversy exists over the role of beta oscillatory activity in successful inhibition.

Considering that gamma oscillations in the motor system are regarded as prokinetic, one would expect them to decrease during successful inhibition. While there is some evidence for decreased gamma-band power (Alegre et al., 2013), most intracranial electrophysiology studies have reported a brief *increase* centered around 70Hz in response to a stop-signal. This phenomenon has been observed in the preSMA, rIFC (Bartoli et al., 2018; Fonken et al., 2016; Swann et al., 2012) and the STN (Fischer et al., 2017; Ray et al., 2012). Generally, this increased gamma activity was present regardless of the success of stopping, but before SSRT (but see Fischer et al., 2017). It is unclear if it reflects an attentional signature for detecting the stop-signal or if it might be involved in the actual implementation of the inhibitory process.

The aforementioned electrophysiological studies are highly informative but correlative in nature. It is therefore not possible to infer whether beta or gamma oscillations are *causally* involved in motor inhibition. Investigation of causal oscillation-function relationships requires experimental control over the strength and/or phase of the ongoing brain rhythms. This can be achieved with transcranial alternating current stimulation (tACS) (Helfrich et al., 2014; Herrmann et al., 2016; Thut et al., 2011). Gamma-band tACS (70Hz) over the primary motor cortex (M1) increases movement amplitude, force development and velocity (Guerra et al., 2018; Joundi et al., 2012; Moisa et al., 2016). Whereas, beta-band stimulation (20 Hz) over M1 results in reduced movement output (Guerra et al., 2018; Joundi et al., 2012; Pogosyan et al., 2009; Wach et al., 2013). Only one previous study assessed the effects of beta- and gamma-band tACS on motor inhibition. Joundi et al. (2012) found that 20Hz tACS over M1 promotes inhibition of unintended movements in the context of a go/no-go task, but 70Hz stimulation did not influence inhibitory performance.

In a go/no-go task either a go cue or a no-go cue is presented, therefore performance on the task likely reflects action restraint, i.e. the decision to respond or not, rather than the ability to inhibit a prepotent response (Leunissen et al., 2017; Raud et al., 2020). In a stop-signal paradigm, stop-signals are presented on a minority of the trials *after* the participant has already begun to initiate their response to the go cue. Thus, stop-signal paradigms are better suited to assess the ability to cancel an already initiated action. In addition, the paradigm lends itself well to models of action cancellation such as the dependent process model (DPM)(Dunovan et al., 2015; Dunovan & Verstynen, 2019). Which can provide insight in whether beta and gamma stimulation affect performance through modulation of the go and/or stop process.

Although go and stop processes ultimately converge upon M1 (Stinear et al., 2009), stopping is triggered upstream of M1. Given the fronto-central topography of beta power during stopping, the goal of the present study was to stimulate preSMA instead of the M1 target used previously. Furthermore, if gamma oscillations observed in the preSMA for successful stops are causally related to the inhibitory process, then gamma stimulation of preSMA might also promote inhibition. To test our hypotheses, we delivered short trains of tACS while participants performed an anticipated response stop-signal paradigm. We hypothesized that beta, and possibly also gamma stimulation targeting preSMA would facilitate response inhibition, i.e. reduce SSRT.

Besides the behavioral consequences of tACS, more direct evidence of the efficiency of tACS in modulating oscillatory brain activity could come from changes in neurophysiological measures. Most evidence so far depends on resting-state measures obtained directly *after* tACS. For example, the after-effects of alpha tACS are thought to rely on plasticity-related changes evoked by spike-timing dependent plasticity (STDP) (Zaehle et al., 2010). While it has been shown that beta and gamma tACS can affect cortical excitability and inhibition (Heise et al., 2016; Nowak et al., 2018; Wischnewski et al., 2019b), the after-effects on spectral power are unknown. Ideally one would be able to assess changes in oscillatory activity *during* tACS, however it remains unresolved whether the tACS artifact can be proficiently removed from concurrent magneto- and electroencephalography (M/EEG) recordings (Neuling et al., 2017; Noury & Siegel, 2018). Here, we opted to use an intermittent tACS design which allows for the comparison between EEG spectral power in trials directly following stimulation with those further removed from the stimulation. This approach avoids the tACS artifact in the EEG yet is less dependent on plasticity related changes. We hypothesized that the increased spectral power due to tACS entrainment might still be visible in the first few seconds after stimulation, but then fade away.

Finally, we performed individual simulations of the electric fields during tACS based on the registered electrode placement and individual MRI scans to assess whether the prospective behavioral effects of the stimulation follow a dose-response relationship.

## Results & Discussion

The main aim of this study was to investigate the role of beta- and gamma-band oscillations in motor inhibition. To this end, we used tACS over preSMA to entrain the 20Hz (beta) and 70Hz (gamma) frequencies. In accordance with previous literature we hypothesized beta stimulation to have an inhibitory effect on motor output during both going and stopping. In contrast, we reasoned that 70Hz stimulation over the preSMA might facilitate movement on go trials, yet also promote inhibition during stop trials. All participants tolerated the stimulation well as indicated by the low ratings of discomfort and fatigue and the ratings did not differ between the two stimulation sessions (Supplementary Table 1).

### Task performance

The independent race model used to estimate SSRT assumes that the distribution of the finishing times of the go process is the same on go and stop trials (context independence). In practice, this means that the average goRT should be higher than the average RT on failed stop trials, i.e. only the fastest go processes are able to escape inhibition. One participant was excluded from all further analyses due to violation of this assumption (Verbruggen et al., 2019).

The number of errors on go trials was matched between stimulation frequencies (F_(1,102)_=0.14, p=0.71), and stimulated versus non-stimulated trials (F_(1,102)_=0.04, p=0.85)(Table 1). The dynamic tracking procedure resulted in a stop success rate close to 50% in both stimulation sessions (no main effect of tACS FREQUENCY: F_(1,102)_=0.8, p=0.36). However, the stop success rate was consistently ~0.5% higher in stimulated than non-stimulated trials (F_(1,102)_=26.8, p<.0001). Besides a real effect of stimulation this could also be caused by the difference in trial numbers between the conditions (40% of all trials were stimulated).

**Table 1.** Effect of stimulation on stop-signal task performance.

| Trial type |  | 20Hz<br>Non-stimulated | 20Hz<br>Stimulated | 70Hz<br>Non-stimulated | 70Hz<br>Stimulated | LME statistics |  |  |  |
| --- | --- | --- | --- | --- | --- | --- | --- | --- | --- |
|  |  |  |  |  |  | df,<br>Error df | Main effect of<br>frequency<br>(20Hz, 70Hz) | Main effect of<br>stimulation<br>(ON, OFF) | Interaction |
| Go | % early response | 0.0 | 0.0 | 0.0 | 0.0 | NA | NA | NA | NA |
|  | % no response | 1.7±2.29 | 1.57±2.46 | 1.56±2.24 | 1.53±2.48 | 1,<br>102 | F=0.14<br>p=0.705 | F=0.04<br>p=0.851 | F=0.16<br>p=0.694 |
|  | goRT (ms) | 17.96±43.64 | 16.90±43.73 | 18.10±42.4 | 17.15±42.6 | 1,<br>28138 | F=0.69<br>p=0.407 | F=3.71<br>p=0.054 | F=0.0<br>p=0.968 |
| Stop | Stop fail RT (ms) | -1.5±34.09 | -2.40±34.14 | -2.47±33.74 | -2.00±33.24 | 1,<br>6937 | F=0.27<br>p=0.606 | F=0.05<br>p=0.818 | F=0.72<br>p=0.395 |
|  | % inhibit | 51.26±0.58 | 51.75±0.69 | 51.38±0.64 | 51.78±0.7 | 1,<br>102 | F=0.8<br>p=0.359 | **F=26.8**<br>**p<.0001** | F=0.3<br>p=0.611 |
|  | SSRT (ms) | 189.4±12.42 | 191.1±11.24 | 193.36±12.22 | 192.4±12.3 | 1,<br>102 | **F=5.36**<br>**p=0.023** | F=0.1<br>p=0.748 | F=1.37<br>p=0.244 |

GoRT and stop fail RT are expressed relative to the target (i.e. response – 800 ms). Mean ± standard deviation is reported. LME = linear mixed model. Results for LME models are given as Type III sums of squares for sequentially fitted fixed effects.

### 20Hz stimulation reduced force production on stop-signal trials

In corroboration with Joundi et al. (2012), the percent change calculations revealed that 20Hz stimulation significantly decreased peak force and peak force rate on successful stop trials by 11.02% and 9.8% respectively (Table 2, Figure 1A,C). Due to the reduction in force output 20Hz stimulation also shortened the time to peak by 2.8% in successful stop trials. In addition, the proportion of successful stop trials with perfect inhibition, i.e. a force trace that remained below 5 times the standard deviation of the baseline period, increased by 4.23% with 20Hz stimulation.

**Figure 1.**
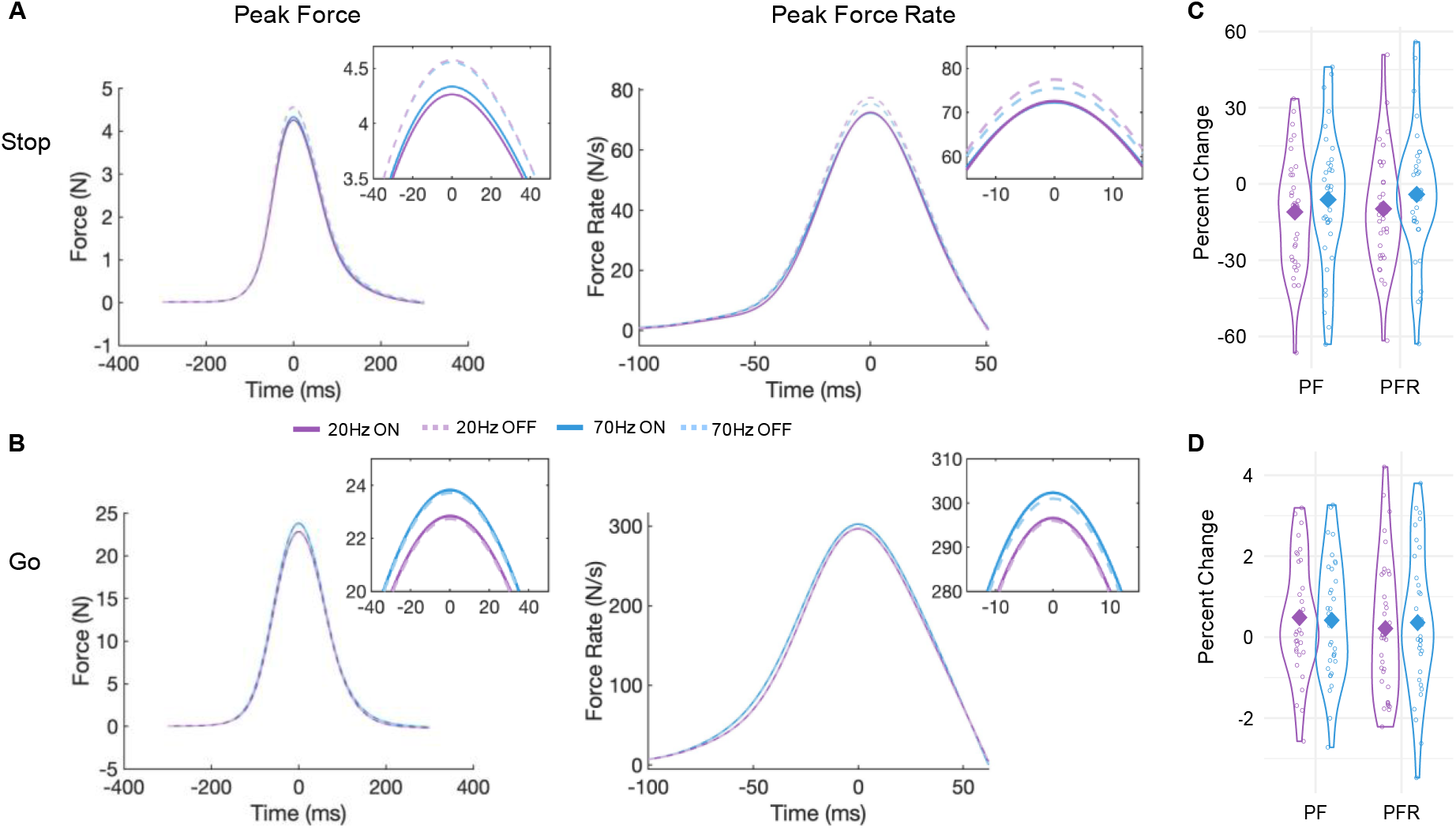
Grand averages for stop (A) and go (B) aligned to peak force and peak force rate for 20Hz (purple) and 70Hz (blue) stimulated (solid lines) and non-stimulated (dashed lines) trials. Sub-windows depict a zoomed view on the peaks. (C-D) Individual percent changes of peak force and peak force rate due to 20Hz (purple) and 70Hz (blue) stimulation on stop (C) and go (D) trials. Solid diamond shape represents the group mean. PF = peak force, PFR = peak force rate.

**Table 2.** Force outcomes on Go and successful Stop trials, with 20Hz or 70Hz stimulation ON or OFF.

|  | 20Hz |  |  |  | 70Hz |  |  |  | LME statistics |  |  |  |  |  |  |  |
| --- | --- | --- | --- | --- | --- | --- | --- | --- | --- | --- | --- | --- | --- | --- | --- | --- |
|  | Go |  | Successful Stop |  | Go |  | Successful Stop |  |  |  |  |  |  |  |  |  |
|  | OFF | ON | OFF | ON | OFF | ON | OFF | ON | df,<br>Error df | tACS<br>FREQ<br>(20Hz,<br>70Hz) | STIM<br>(ON,OFF) | TRIAL<br>TYPE<br>{succ<br>stop,go) | tACS<br>FREQ*<br>STIM | tACS<br>FREQ*<br>TYPE | STIM*<br>TYPE | tACS<br>FREQ*<br>STIM*<br>TYPE |
| Peak force<br>(N) | 22.87<br>±7.81 | 22.97<br>±7.85 | 4.68<br>±4.21 | 4.38<br>±4.29 | 24.04<br>±7.47 | 24.17<br>±7.48 | 4.56<br>±4.33 | 4.42<br>±4.31 | 1,<br>34353 | **F=186.24**<br>**p<.0001** | F=0.01<br>p=0.908 | **F=68939.6**<br>**p<.0001** | F=0.19<br>p=0.663 | **F=62.76**<br>**p<.0001** | **F=6.45**<br>**p=0.011** | F=0.61<br>p=0.436 |
| %change | **0.48±1.41**<br>**t=2.04, p=0.05** |  | **-11.02±22.23**<br>**t=-2.93, p=0.006** |  | 0.42±1.45<br>t=1.7, p=0.1 |  | -6.17±26.55<br>t=-1.37, p=0.18 |  |  |  |  |  |  |  |  |  |
| Peak force<br>rate (N/s) | 297.42<br>±104.1 | 289.56<br>±104.9 | 79.92<br>±67.62 | 75.36<br>±69.29 | 305.26<br>±98.23 | 307.12<br>±99.1 | 77.36<br>±70.88 | 75.59<br>±70.03 | 1,<br>34289 | **F=25.97**<br>**p<.0001** | F=0.04<br>p=0.848 | **F=46851.9**<br>**p<.0001** | F=0.87<br>p=0.351<br>3 | **F=16.19**<br>**p<0.001** | **F=6.27**<br>**p=0.012** | F=0.55<br>p=0.458 |
| %change | 0.21±1.68<br>t=0.75, p=0.46 |  | **-9.8±22.23**<br>**t=-2.61, p=0.014** |  | 0.36±1.73<br>t=1.23, p=0.23 |  | -4.13±25.25<br>t=-0.97, p=0.34 |  |  |  |  |  |  |  |  |  |
| Time to<br>peak (ms) | 147.75<br>±47.57 | 148.17<br>±48.37 | 99.63<br>±50.94 | 98.07<br>±50.46 | 150.79<br>±47.01 | 151.04<br>±47.77 | 100.46<br>±51.95 | 99.45<br>±51.91 | 1,<br>33329 | **F=36.41**<br>**p<0.001** | F=0.346<br>p=0.556 | **F=7501.43**<br>**p<0.001** | F=0.058<br>p=0.809 | F=3.310<br>p=0.069 | F=2.909<br>p=0.088 | F=0.222<br>p=0.638 |
| %change | 0.47±2.17<br>t=1.27, p=0.21 |  | **-2.77±7.18**<br>**t=-2.28, p=0.03** |  | 0.003±2.8<br>t=0.0, p=0.99 |  | -2.53±12.88<br>t=-1.16, p=0.25 |  |  |  |  |  |  |  |  |  |
| Proportion<br>of trials with<br>force >5*<br>baseline SD |  |  | 0.81<br>±0.12 | 0.77<br>±0.13 |  |  | 0.81<br>±0.11 | 0.78<br>±0.15 | 1,<br>102 |  |  | F=0.062<br>p=0.804 | **F=4.707**<br>**p=0.032** |  | F=0.161<br>p=0.689 |  |
| %change |  |  | **-4.23±9.66**<br>**t=-2.59, p=0.014** |  |  |  | -3.25±10.92<br>t=-1.76, p=0.09 |  |  |  |  |  |  |  |  |  |

3 Mean ± standard deviation is reported. LME = linear mixed model. Results for LME models are given as Type III sums of squares for sequentially fitted fixed effect

On go trials, 20Hz stimulation did not affect peak force rate or the time to peak but did result in an 0.48% increase in mean peak force (Table 2, Figure 1B,D). This might seem counterintuitive and contradicts the findings from Joundi et al. (2012). However, it has been demonstrated before that the response force on go trials increases with the increasing likelihood of a stop-signal appearing, in other words when the readiness to respond is low (van den Wildenberg et al., 2003). Analogous to the relationship between excessive beta oscillation and the bradykinesia and rigidity symptoms in PD (Kuhn et al., 2006), we speculate that 20Hz stimulation puts a global break on the system, making it harder to respond when necessary. Therein also lies an important difference between the go/no-go task and our anticipated response stop-signal paradigm. In the go/no-go task there is no hard constraint on when to respond other than the experimenter’s instruction to respond as fast as possible. As a result, participants tend to slow down until they gather enough evidence for going (Leunissen et al., 2017; Szmalec et al., 2009). Here, participants needed to perform a response at a known point in time, and on top of that there was visual feedback on their performance, reinforcing go task performance.

### Higher force production on go trials in the 70Hz stimulation session

Peak force and peak force rate were significantly higher in the 70Hz than in the 20Hz stimulation session (Table 2, Figure 1B). This effect seemed to be driven by the go trials (significant TRIAL TYPE* tACS FREQUENCY interaction; estimated difference in peak force on go trials: 1.06±0.06N, z=16.53, *P*_adjusted_<.0001, stop trials: −0.09±0.13N, z=-0.691, *P*_adjusted_=0.896; estimated difference in mean peak force rate on go trials: 6.12±0.92N/s, z=6.68, *p*_adjusted_<.0001, stop trials: −2.23±1.86N/s, z=-1.2, *P*_adjusted_=0.617). The percent change calculations show that peak force on stimulated go trials in the 70Hz stimulation session was 0.42% higher than peak force on the non-stimulated go trials. This increase was not significant however (*p*=0.1). Since there was no effect of stimulation ON/OFF or a separate sham session it is impossible to infer whether the difference in force on go trials between the 20Hz and 70Hz session is caused by a decrease due to 20Hz stimulation or an increase due to 70Hz stimulation. Based on the findings from Joundi et al. (2012), the most likely scenario is perhaps a combination of both.

The lack of differences between stimulated and non-stimulated go trials in the 70Hz stimulation session could be caused by several factors. First, the go task involves a timed response rather than a reaction to an external cue. It is possible that gamma oscillations are less involved in timed responses. Second, the preSMA and SMA-proper are thought to be responsible for linking situations with appropriate actions (Nachev et al., 2008). Hosaka et al. (2016) demonstrated that gamma oscillatory activity in the (pre)SMA of monkeys increased during movement, but particularly when the action plan needed to be updated. Finally, given the significant difference in peak force and peak force rate between the 20Hz and 70Hz stimulation session it is also possible that the stimulation effects carried over to the non-stimulated trials (see tACS aftereffects section for a more in-depth discussion).

### Opposing effects of 20Hz and 70Hz stimulation on braking drift rate

No difference was observed in goRT between the different stimulation frequencies or stimulation ON/OFF, although the latter showed a trend towards shorter response times (i.e. closer to the target) with stimulation ON in both the 20Hz and 70Hz stimulation session (F_(1,28138)_=3.71, *p*=0.054)(Table 1). This might be related to the increased force and velocity observed in go trials during stimulation, as discussed above. The absence of a stimulation effect on goRT corroborates the findings of Pogosyan et al. (2009) and Joundi et al. (2012).

SSRT was significantly shorter in the 20Hz than in the 70Hz stimulation session (F_(1,102)_=5.36, p=0.023), but there was no difference between SSRT estimated from stimulated and non-stimulated trials, or a FREQUENCY*STIMULATION interaction. Again, this precludes us from concluding whether 70Hz stimulation increased SSRT or 20Hz stimulation decreased SSRT.

SSRT is an estimate of the covert latency of the stop signal estimated based on the independent race model (Logan et al., 1984). Conceptualizing the go and stop processes as independent processes racing against each other has accounted well for the observed behavioral data in the stop-signal paradigm (Matzke, 2018). However, on a neural level it is evident that the neurons involved in movement initiation and inhibition interact with each other during action cancellation (Boucher et al., 2007; Munoz & Schall, 2003; Schmidt et al., 2013). Behavioral models that include such a dependency between the go and stop process indeed provide an even better fit to the data (Boucher et al., 2007; Dunovan et al., 2015). In the dependent process model (DPM) from Dunovan et al. (2015; 2019) the go process is modelled as a stochastic accumulator that gathers evidence at a certain driftrate (*V_e_*), leading to a response when it crosses an upper threshold (*a*) (Figure 2).

**Figure 2.**
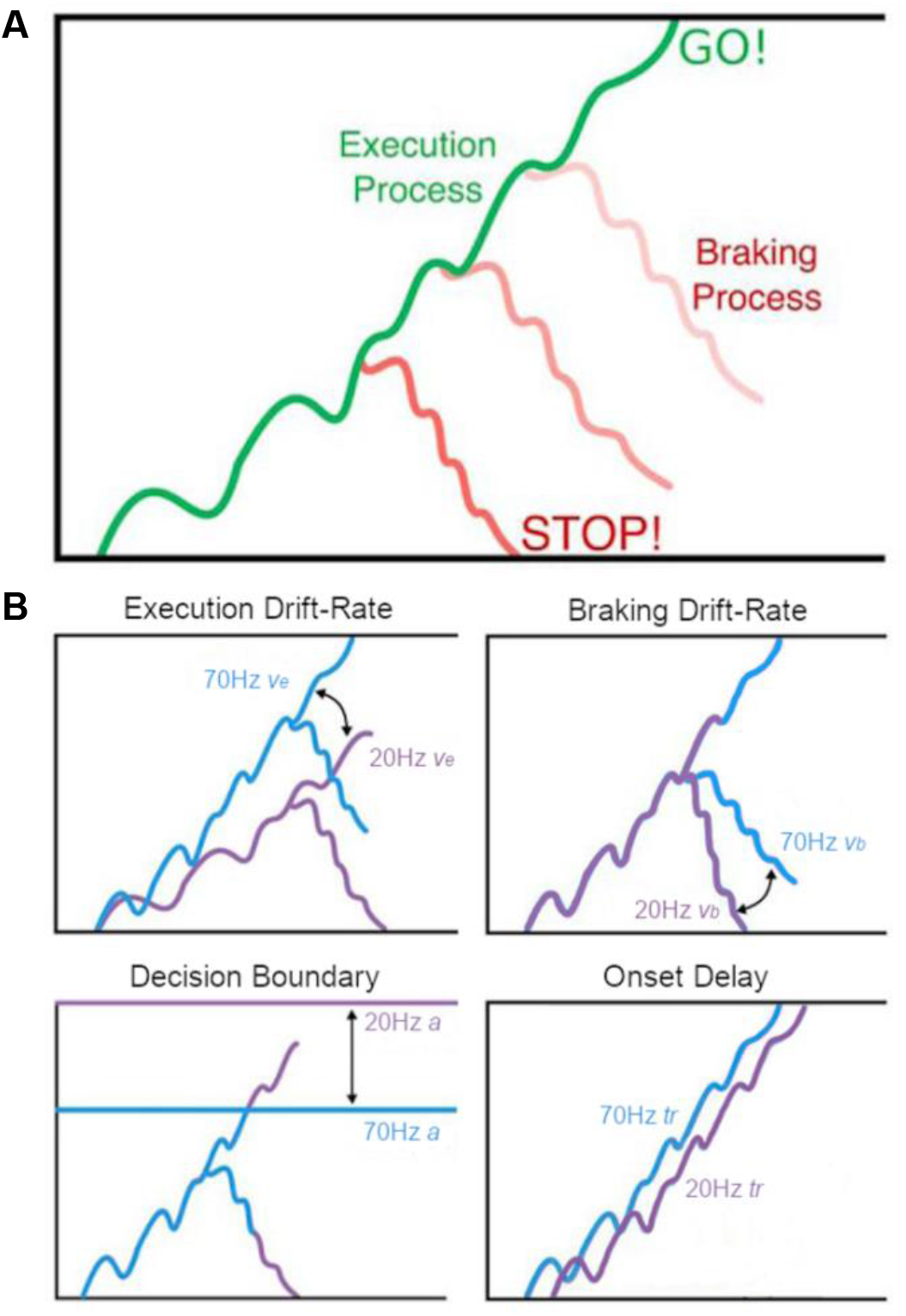
Graphical description of the dependent process model (DPM). (A) The DPM assumes that the state of an accumulating execution process at the time of the stop-signal determines the initial state of the braking process, making it more difficult to cancel actions closer to the execution boundary. (B) Possible control mechanisms that could be altered by beta (20Hz) and/or gamma (70Hz) tACS stimulation. Adapted with permission from Dunovan and Verstynen (2019).

In the event of a stop-signal, a second braking process is instantiated at the current state of the execution process and must reach the bottom boundary before the execution threshold is reached in order to cancel motor output. This model not only provides a better fit to the data, but also gives insight into the mechanisms underlying going and stopping. Another advantage is that the DPM takes into account the full goRT and failed stop RT distributions. Even though there was no difference in average goRT and SSRT between stimulated and non-stimulated trials, stimulation might have altered the shape of the response distributions. By fitting the DPM differences in shape can be picked up and are reflected in a change in the rate of the execution (*v_e_*) or braking drift (*v_b_*), shift the onset time at which the execution process begins to accumulate (*tr*), or change the distance to the threshold (*a*).

The braking drift modulation model best explained the effect of stimulation on task performance (Figure 3, Table 3). Braking drift rate increased due to 20Hz stimulation, whereas 70Hz stimulation decreased the braking drift rate (note that more negative values reflect a stronger braking process). Although the braking drift modulation model provided the best fit to the data the other models also had very good fits, suggesting that stimulation might not solely affect braking drift.

**Figure 3.**
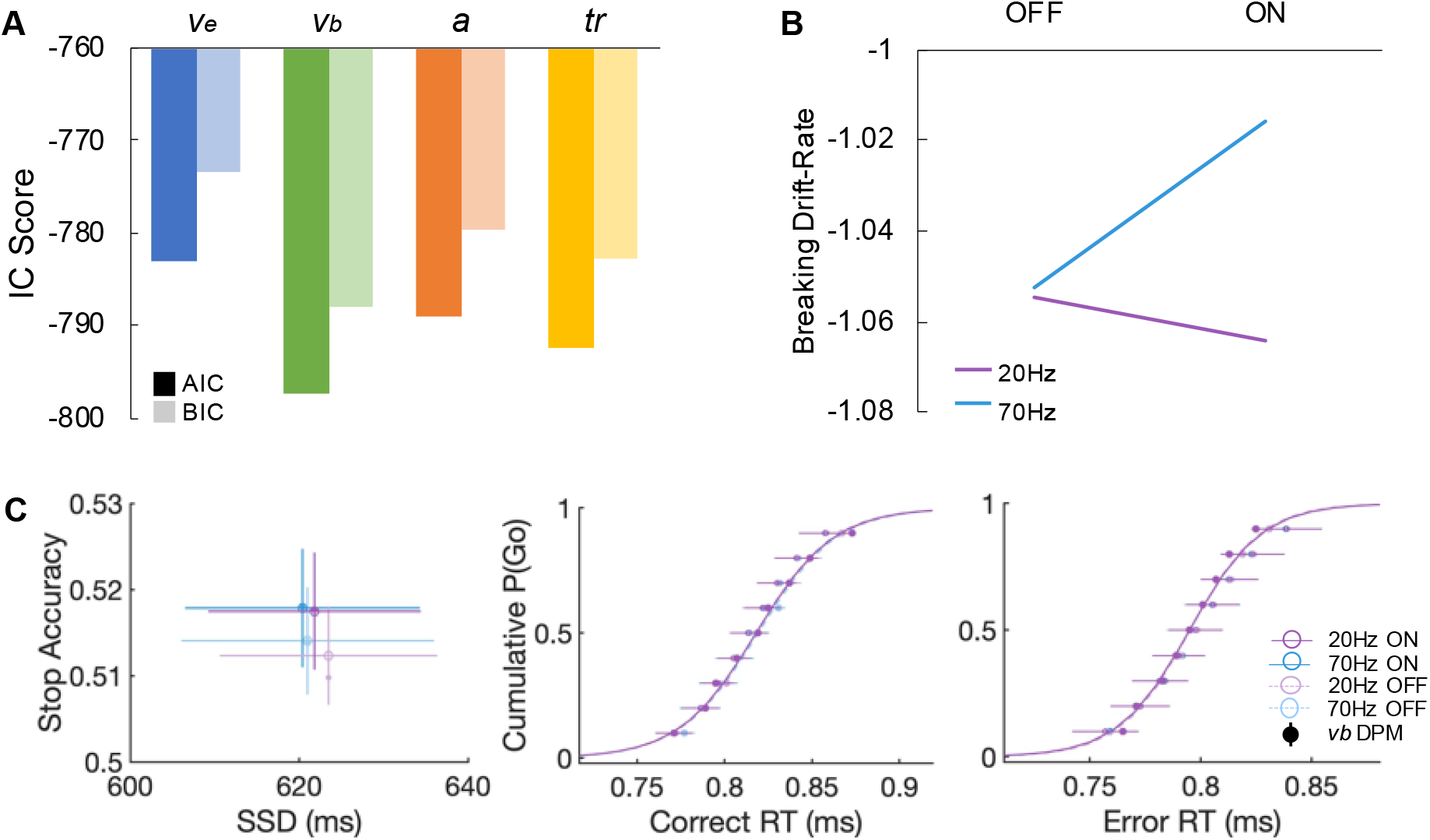
(A) Goodness-of-fit measures for the four different dependent process models. AIC (dark) and BIC (light) scores for all single-parameter models, allowing either execution drift-rate (*V_e_*; blue), braking drift-rate (*V_b_*; green), execution boundary height (*a*; orange), or onset delay (*tr*; yellow) to vary across conditions. The model with the lowest score, in this case the braking drift modulation model, is preferred. (B) Parameter estimates of the braking drift rate in stimulated and non-stimulated trials in the beta (20Hz) and gamma (70Hz) sessions. (C) Model predicted data (solid lines and circles) simulated with best-fit parameters from the *V_b_* model overlaid on the average ± SEM empirical data (transparent circles and horizontal lines).

**Table 3.**
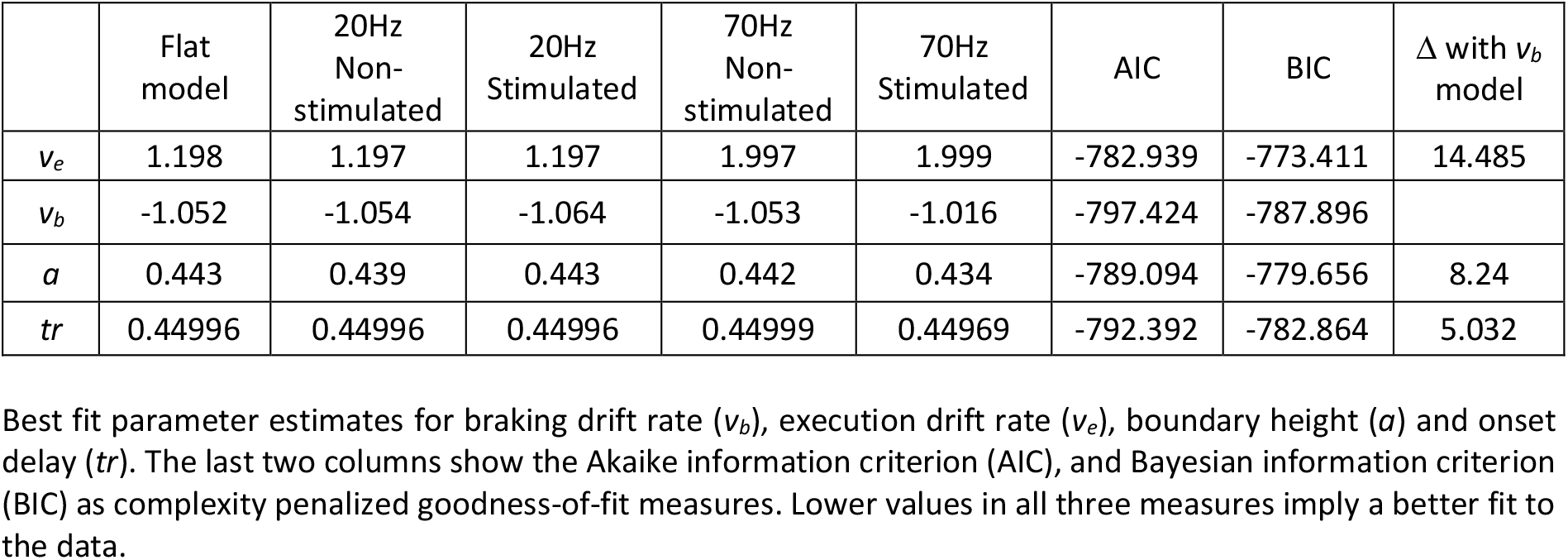
Dependent process model parameter estimates and fit statistics.

| | Flat model | 20Hz Non-stimulated | 20Hz Stimulated | 70Hz Non-stimulated | 70Hz Stimulated | AIC | BIC | $\Delta$ with $v_b$ model |
| --- | --- | --- | --- | --- | --- | --- | --- | --- |
| $v_e$ | 1.198 | 1.197 | 1.197 | 1.997 | 1.999 | -782.939 | -773.411 | 14.485 |
| $v_b$ | -1.052 | -1.054 | -1.064 | -1.053 | -1.016 | -797.424 | -787.896 | |
| $a$ | 0.443 | 0.439 | 0.443 | 0.442 | 0.434 | -789.094 | -779.656 | 8.24 |
| $tr$ | 0.44996 | 0.44996 | 0.44996 | 0.44999 | 0.44969 | -792.392 | -782.864 | 5.032 |

Best fit parameter estimates for braking drift rate ( $v_b$ ), execution drift rate ( $v_e$ ), boundary height ( $a$ ) and onset delay ( $tr$ ). The last two columns show the Akaike information criterion (AIC), and Bayesian information criterion (BIC) as complexity penalized goodness-of-fit measures. Lower values in all three measures imply a better fit to the data.

Taken together, the results from the force and response time analyses show opposing roles for beta and gamma oscillations that fit with the prevailing view that gamma activity in the motor system is pro-kinetic, while beta oscillations support motor suppression. Stimulation interacts with the beta rhythm to drive oscillations, but the degree to which this resonance phenomenon takes place is dynamically determined by task demands, e.g. 20Hz tACS had a much stronger effect on stop-signal trials than go trials.

We did not find evidence for a causal role of gamma oscillations in stopping. Gamma stimulation even seemed to reduce the speed of the braking process, and rather affected go performance. Peak force and peak force rate were higher in the 70Hz stimulation session. Moreover, the parameter estimates of the DPM’s all point towards movement facilitation with 70Hz stimulation (i.e. increased execution drift rate, lower boundary heights and shorter onset delay). Fischer et al. (2017) suggested that comparisons between executed and withheld movements might reflect the lack of movement rather than the stopping process per se. By using a task in which continuous ongoing movement needs to be inhibited they circumvented this issue and, based on their findings, they advocate that increased gamma and not beta activity is responsible for successful inhibition. We want to emphasize that our paradigm was very successful in ensuring go response initiation, since the proportion of successful stop trials in which we could still identify a force response 5 SD above baseline was on average 80% (range 50-100%, Table 2) opposed to ~45% in Joundi et al. (2012). Therefore, we find this explanation unlikely for our findings.

Gamma activity is thought to reflect local activity, whereas beta band activity seems important for long-distance communication between frontal cortex and the basal ganglia (Bartoli et al., 2018; Swann et al., 2012). This long-distance communication might require less precise timing of the entrainment. Because preSMA gamma-band activity increases for both going and stopping (albeit at different timescales), stimulating at 70Hz for the whole trial duration might create a conflict between facilitating movement versus promoting inhibition.

### Sources of variability

The behavioral results largely follow the hypothesized effects of stimulation. However, it is also evident that the effects are variable from one participant to the next. In an attempt to identify some possible sources of this variability we performed several explorative analyses.

#### Electrical field modeling

The amount of current that reaches the targeted brain area likely influences the size of the stimulation effect. Stimulation was provided at a fixed output current of 1mA, but individual differences (e.g. in skull thickness and scalp to cortex distance) can influence how much current actually reaches the brain (Datta et al., 2012). To investigate whether there was a dose-response relationship between amount of current reaching the preSMA and the behavioral effect of stimulation we modelled the current flow in each individual based on the registered electrode positions and MRI scans. Two recent studies provide validation for the accuracy of such models by using intracranial recordings (Huang et al., 2017; Opitz et al., 2016).

The modeling results indicated that, on average, the current successfully reached the preSMA and that the field expansion was limited to the area between the four return electrodes (Figure 4A). The predicted normalized electrical field strength in the preSMA ROI during the beta session was significantly related to the percent change in peak force on successful stops (r=-0.469, p=0.028) (Figure 4B). This dose-response relationship supports the notion that tACS stimulation has a causal effect on behavior, and suggests that it would be advisable to try to control the amount of current to the brain by adjusting the output current based on the current flow predictions (Bestmann & Ward, 2017; Tan et al., 2020).

**Figure 4.**
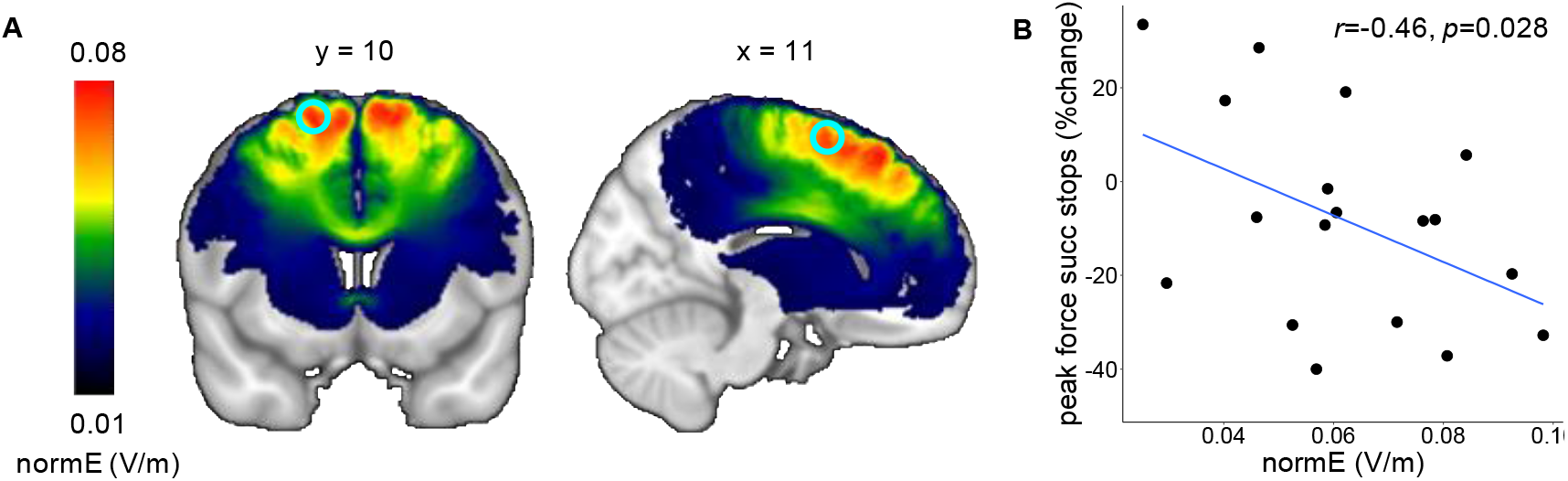
Simulation of the electrical field of the tACS. A) Group average of the normalized predicted electrical field distribution in MNI space. The cyan circle indicates the preSMA ROI (10mm sphere around coordinate [11,10,62] based on a previous fMRI study with the same task paradigm). B) Relation between normalized predicted electrical field strength within the preSMA ROI and the effect of beta stimulation on peak force in successful stop trials. The shaded area represents the 95% confidence interval.

#### Individual peak frequency

Another possible source of variability is the stimulation frequency. The effects of tACS seem to follow an Arnold tongue principle in the sense that tACS can only modulate ongoing brain oscillations if the frequency of the tACS is very close to the frequency of the intrinsic brain oscillations. To be able to synchronize or entrain frequencies further away from the “Eigenfrequency” the external driving force (tACS) will need to be stronger (i.e. higher stimulation amplitude) (Ali et al., 2013). In this study we chose to use 20Hz and 70Hz as stimulation frequencies because oscillatory activity in the motor system is commonly centered around these frequencies (Chakarov et al., 2009; Fischer et al., 2017; Muthukumaraswamy, 2010).

To evaluate whether individuals with a peak in beta oscillatory power close to 20Hz responded more strongly to the stimulation than individuals with a peak frequency further removed from 20Hz, we identified the individual beta frequency based on the resting state EEG acquired before the beta stimulation session and plotted it against the percent change in force on successful stop trials. Figure 5 demonstrates that participants with a peak between 18-22Hz typically showed a decrease in peak force on successful stops with beta stimulation for 23/27 participants (85%), whereas outside of that range 3/8 participants (37%) showed a decrease. This corroborates with findings from Vossen et al. (2015) who found the after effects of alpha stimulation to be the strongest at the individual alpha peak and not present ± 2Hz away from the individual peak frequency. Note that a similar procedure is not possible for the gamma-band as the signal to noise ratio with scalp EEG makes it difficult to reliably uncover the higher frequencies and gamma-band activity is typically quite broad without a clear individual peak.

**Figure 5.**
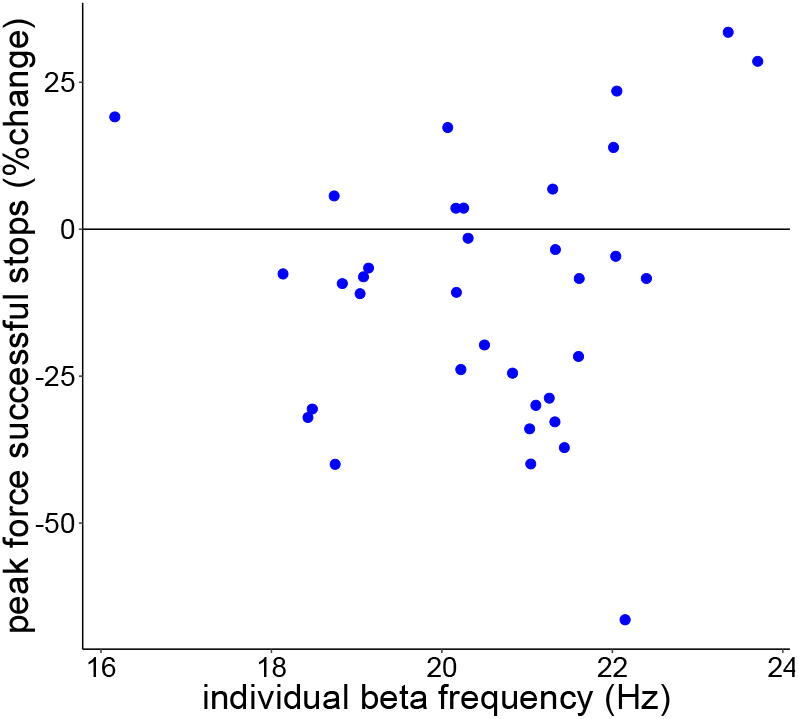
Estimated individual beta peaks based on pre-stimulation resting EEG at electrode Fz plotted against percent change in peak force on successful stop trials with 20Hz stimulation. Participants that fall within the grey shaded area (i.e. within 2Hz from the stimulation frequency) tend to show improved inhibitory performance during 20Hz stimulation.

#### tACS aftereffects

The central tACS electrode was placed under channel FCz, therefore we focused our analyses on channel Fz, which lies directly in front of FCz and still covers the preSMA. To give further justification for this choice, we contrasted all non-stimulated successful stop and go trials from both sessions with each other. This comparison revealed one significant positive cluster (*p*=0.0002) with a fronto-central topography in which the beta activity was higher for successful stop trials than in go trials from 150 till 400ms after the presentation of the stop signal. This cluster includes electrode Fz and extended to electrodes covering the rIFC (Figure 6A). Moreover, participants with higher Fz beta activity 150ms after the stop signal had shorter SSRTs (average non-stimulated SSRT over both sessions, *r*=-0.44, *p_adjustedr_*=0. 031)(Figure 6B).

**Figure 6.**
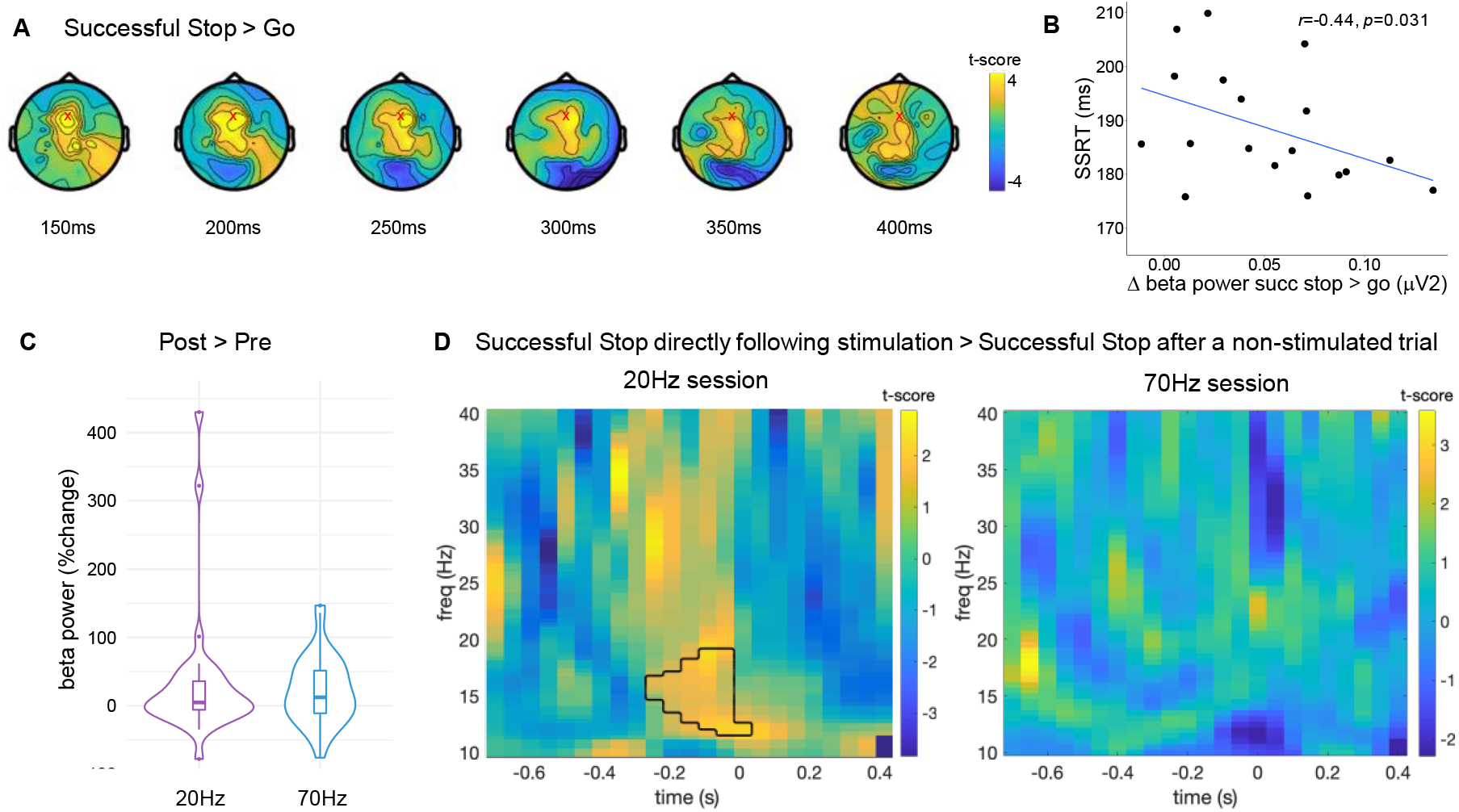
(A) Topographic distribution of increased beta activity in successful stops versus go trials 150-400ms after the presentation of the stop signal. The red X marks the location of electrode Fz. (B) Correlation between the difference in beta power for successful stop and go trials and stop-signal reaction time (SSRT) in electrode Fz. The shaded area represents the 95% confidence interval. (C) Percent change in resting EEG beta power (13-30Hz) from pre to post stimulation. (D) Time-frequency power plots of the comparison between successful stop trials that directly followed stimulation and successful stop trials that occurred after a non-stimulated trial. The black outline represents the significant positive cluster.

#### After-effect of beta stimulation is state-dependent

tACS effects have been shown to outlast the stimulation duration (Veniero et al., 2015). These after-effects are thought to rely on changes occurring through spike-timing dependent plasticity (Veniero et al., 2015; Zaehle et al., 2010), rather than the entrainment of the oscillations which takes place online (Helfrich et al., 2014). For 20Hz tACS, after-effects on cortical excitability have been found to last up to an hour (Wischnewski et al., 2019a). Here, beta oscillatory power at rest was enhanced after 20Hz tACS (pre-versus post-test: V=418, p=0.039). However, this effect was also present in the 70Hz stimulation condition (V=467, p=0.012), and the change in beta power from pre-to post-stimulation did not differ between the two stimulation frequencies (V=276, p=0.723) (Figure 6C). This could mean that the increase in beta power is due to general processes such as performing the task, or that 70Hz stimulation influenced beta power (de Hemptinne et al., 2013).

Comparing successful stop trials directly following stimulation (i.e. stop-signal was presented ~2.5s after the end of the previous stimulation train) with successful stop trials following a non-stimulated trial (i.e. stop-signal was presented ~7s after the of end of the previous stimulation train) revealed one significant positive cluster with higher beta activity 250-50ms before the presentation of the stop signal in trials directly following 20Hz stimulation (Figure 6D). This effect was not present in the 70Hz stimulation session. Under normal circumstances beta activity significantly increases after the presentation of the stop signal (Figure 6A)(Swann et al., 2012; Wagner et al., 2018; Wessel et al., 2016). The finding that stopping was successful when beta activity in preSMA was already increased 200ms before the stop-signal (i.e. before participants knew they had to stop) suggests that beta stimulation increases proactive inhibition. The fact that beta activity was not enhanced during the entire period, but only around the time that a stop-signal could be expected, highlights that the (after) effects of tACS are state-dependent. These results also illustrate that the effects of our intermittent stimulation protocol carried over to the non-stimulated trials and likely clouded the differences between stimulation ON/OFF, similar to the offline effects reported in Heise et al. (2019).

### Conclusion

We provide evidence that fronto-central beta oscillatory activity is causal to stopping ability. During successful stop trials 20Hz stimulation over preSMA resulted in a considerable decrease in force output and the response time models revealed that 20Hz stimulation specifically increased braking drift. These effects followed a dose-response relationship with the strength of the individually simulated electric field. In contrast, 70Hz stimulation seemed to lead to a decrease in braking drift and to mainly affect go task performance. Our results highlight the state-dependency of tACS entrainment and, along with recent complementary research (Sundby et al., 2020), pave the way for the use of fronto-central beta activity as a functional marker of motor inhibition.

## Methods

### Participants

Thirty-six right-handed (laterality quotient range 33-100, mean 90.5 (Oldfield, 1971)) healthy volunteers (age range 19-28y, mean 22.5y, 15 male) were included in this study. Standard screening verified that none of the participants presented with contraindications regarding non-invasive brain stimulation (Bikson et al., 2009; Woods et al., 2016). All procedures were approved by the ethical committee of the University Medical Center of the KU Leuven (protocol no. 57640) and written informed consent was obtained from all participants.

### Experimental design

Participants underwent two tACS-EEG sessions in which they received either 20Hz (beta) or 70Hz (gamma) stimulation during the performance of a stop-signal task. The stimulation frequency order was counterbalanced across participants, and sessions took place at least 48h apart (range 2-55 days, mean 10 days) to avoid potential carry-over effects. Data acquisition in each session started with 5 min of resting EEG with eyes open while fixating on a white fixation cross on a black background (pre-EEG), and also ended with 3 min of resting EEG (post-EEG).

### Stop-signal paradigm

Participants performed an anticipated response stop-signal task (Coxon et al., 2006; Leunissen et al., 2017; Slater-Hammel, 1960). They were comfortably seated at approximately 1m distance of a computer screen (refresh rate 60Hz). The visual display consisted of a vertical indicator, presented centrally on the screen, that moved from the bottom upwards on each trial (Figure 7A). A target line was situated 800ms from onset. The primary task was to stop the indicator at the target by pinching a force transducer (OMEGA Engineering, Norwalk, CT, USA) held between the index finger and thumb of the right hand (go trials). In line with Pogosyan et al. (2009) and Joundi et al. (2012) response force measures were taken, as the more detailed force kinematics were more sensitive to changes in behavior due to tACS stimulation than response times. Participants were instructed to perform these go trials as accurately and consistently as possible. To reinforce go task performance the color of the target line changed to green, yellow, orange or red at the end of each trial, depending on whether responses were within 20, 40, 60, or >60ms of the target. In 33% of the trials the indicator stopped automatically prior to the target. When this happened, participants tried to prevent pressing the sensor (stop trials, Figure 7A). Separate staircasing algorithms were used for stimulated and non-stimulated trials to ensure convergence to 50% success on stop trials in each condition. The initial stop time was set at 250ms from the target and was adjusted in steps of 25ms. The indicator was reset to empty after 1s. The inter-trial interval was 4.5s.

**Figure 7.**
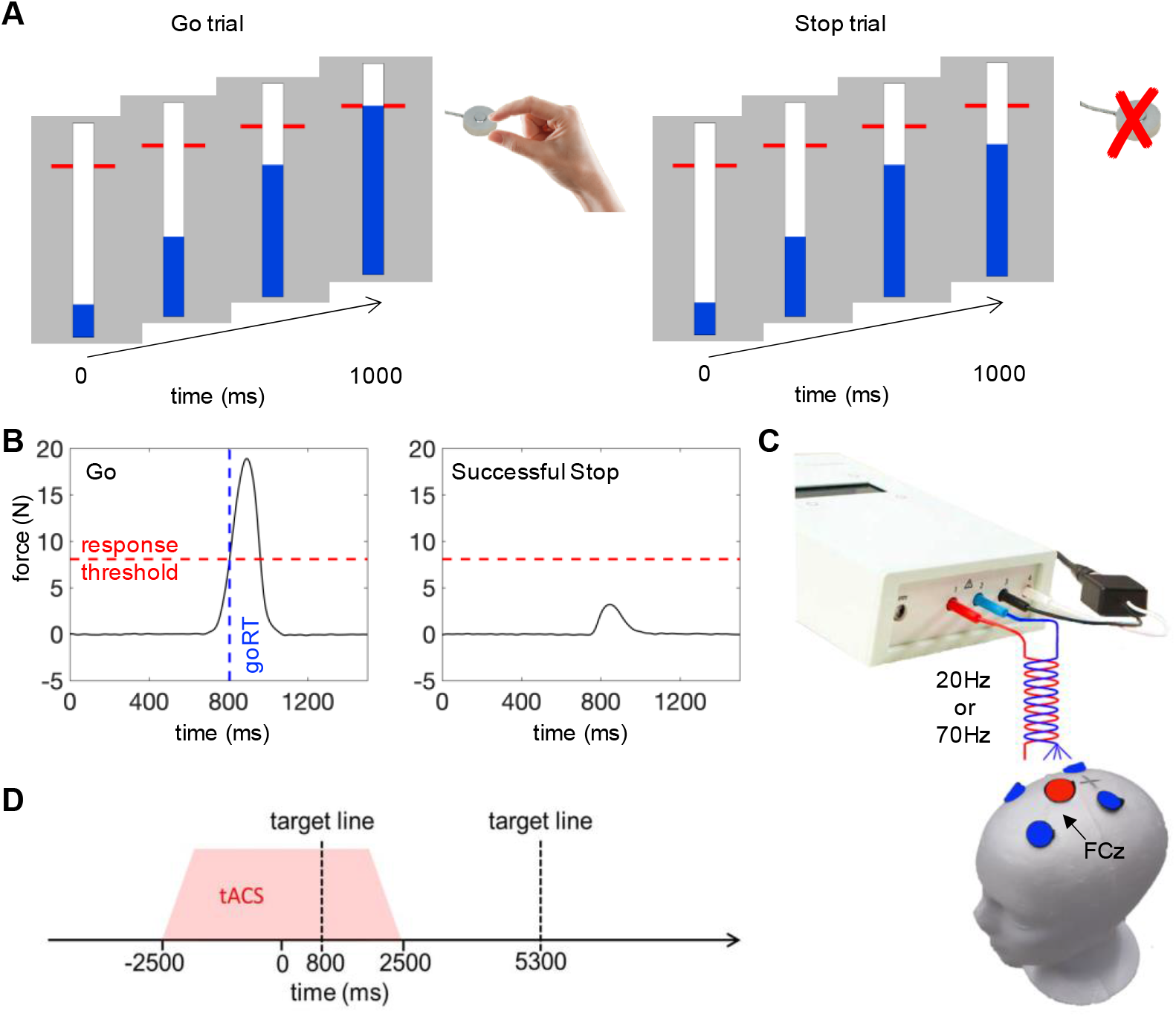
(A) Anticipated response version of stop-signal paradigm. An indicator (depicted in blue) increased from the bottom up at constant velocity reaching the top in 1 s. In ‘go’ trials, participants had to stop the indicator as close as possible to the red target line by squeezing a force sensor. In ‘stop’ trials, the bar would stop filling before it reached the target line and participants were instructed to withhold their response. (B) Example force trace of a go trial and a successful stop with a partial response. Response times were recorded as the time between indicator fill onset and the moment the force signal first exceeded the response threshold (~30% of maximum voluntary force). Stop trials were classified as failed stop trials if the force produced exceeded the response threshold. If the force remained below the threshold, the trial was classified as a successfully inhibited. (C) Electrode montage with center electrode (∅ 2.5cm) over FCz and surrounding electrodes at ~5cm center-to-center distance (∅ 2cm). (D) Event-related alternating current stimulation ensued randomly in 40% of the trials. Stimulation commenced 2.5s before indicator fill onset and lasted for a total of 5s including fading in/out phase of 0.5s. Between the end of the previous and the start of the next stimulation trains was a minimum interval of 4.5s.

Before administration of the experimental task, participants were instructed to pinch the force transducer as hard as possible for ~5 seconds to determine maximal voluntary force (MVF) (custom LabVIEW software, National Instruments, Austin, TX, USA). This procedure was repeated 3 times, and the highest peak-force value was recorded. Next, participants were asked to pinch with a short but powerful pulse against the force transducer, as if they were responding to a stimulus. The response threshold was initially set to 35% of their MVF and lowered in steps of 5% till participants reported they could comfortably cross the threshold (range 20-35%, mean 29%) to avoid fatigue. The force signal was sampled at 1000Hz on each trial from the moment that the indicator started filling for 1.5 seconds. Response times were recorded as the time between indicator fill onset and the moment the force signal first exceeded the threshold. Stop trials were classified as failed stop trials if the force produced exceeded the response threshold. If the force remained below the threshold, the trial was classified as a successfully inhibited (Figure 7B). Participants practiced the task by performing 20 go trials, followed by 20 trials in which go and stop trials were mixed. Participants completed six concurrent tACS-EEG task runs per session, each comprising 67 go and 34 stop trials presented in a pseudorandomized order (606 trials per session in total)

### tACS-EEG procedures

EEG was recorded by means of an EGI 400 Geodesic system with a 128-channel HydroCel Geodesic Sensor Net (EGI, Eugene, OR, USA) and a sampling rate of 1000Hz (Net Station v5.1.2). Cz was used as physical reference during recording and impedance of all electrodes was kept below 50kΩ as recommended for this system. The position of the electrodes on the participants scalp were localized with the Geodesic Photogrammetry System (GPS 2.0, EGI, Eugene, OR, USA).

20Hz (beta) and 70Hz (gamma) tACS were applied in separate sessions using an 4×1 HD-tACS setup (DC Stimulator Plus, NeuroConn, Ilmenau, Germany) with a stimulation intensity of 1000μA (peak-to-peak amplitude). The target electrode (2.5cm Ø) was placed over the preSMA (FCz) (Homan et al., 1987), and the four surrounding electrodes (2cm Ø) were placed at positions F1, F2, C1 and C2 (Figure 7C). In all instances impedance of the tACS electrodes was kept below 10kΩ (range 1.2-7kΩ, mean 3.51kΩ). tACS was applied in an event-related manner, distributed pseudo-randomly over 40% of both go and stop trials. Each stimulation train ramped up in 0.5s, 2.5s before the start of the trial and lasted a total of 4s before ramping down again (Figure 7D). Between the end and the start of the next stimulation train was an interval of 4.5s or 9s. Over one experimental session the participants received a total of 18 min of tACS.

For the first 16 participants, the classic HydroCel Geodesic Sensor Net was used in which the electrodes are encased in plastic cups covered with sponges that are soaked in electrolyte solution. To ensure good contact of the tACS electrodes a sponge soaked in the same electrolyte solution was placed under the rubber electrodes. With this set-up the tACS stimulation caused saturation of several EEG electrodes in about 1/3 of the participants, resulting in large artefacts even during the non-stimulated periods. The remaining 20 participants were tested using HydroCel Geodesic Sensor Net 130 LTM nets, where the cups were filled with electrolyte gel (Redux®, Parker Laboratories, Fairfield, NJ, USA), which resolved the saturation problems. The same gel was used to ensure a good contact between skin and tACS electrodes.

### Evaluation of subjective level of discomfort caused by tACS and self-perceived level of fatigue

The level of discomfort was assessed after each session according to a Visual Analogue Scale (VAS) of 10cm length without numerical indication, extremes constituted ‘absolutely no discomfort/pain’ and ‘worst discomfort/pain ever’. The point on the scale marked by the participant was subsequently converted into a score ranging from 1-10 (Huskisson, 1974). Similarly, participants evaluated their perceived level of fatigue with a VAS (ranging from ‘absolutely not tired’ to ‘maximally tired/exhausted’) at the beginning and end of each experimental session.

### Behavioral analysis

Force data and response times were analyzed using Matlab R2016a (Mathworks, Natick, MA, USA).

#### Force data

Force data was filtered with a fifth-order 20Hz low pass Butterworth filter, and baseline corrected by subtracting the average force between −650 to −300ms prior to the target. Per trial we determined: (i) peak force (i.e. maximum force in that trial), (ii) peak rate of force development, and (iii) the time to peak, which is defined as the time between the first instance that the force trace exceeds 5*SD of the baseline period and the peak. On successful stop trials force production did not always exceed 5*SD of the baseline period. To quantify this and to capture possible changes in the proportion of successful stop trials with ‘perfect inhibition’, we calculated the proportion of successful stop trials in which force production did exceed the threshold. Trials with extreme early responses (>400ms before the target) and go trials where there was no response were considered errors, and trials with force output more than 2.5*SD from their respective mean were defined as outliers and removed. In addition to the average per trial type and condition, we also calculated the percent change in peak force, peak force rate, time to peak, and the proportion of successful stop trials in which the response exceeded 5*SD of the baseline period between stimulated versus non-stimulated trials.

#### Response times

Go trial response times (goRT) and response times for unsuccessful stop trials were determined relative to the target (time of response - 800ms). Early response times (>400ms before the target) and go trials where there was no response were considered errors and removed. For stop trials, the probability of responding was calculated and stop signal reaction time (SSRT) was determined via the integration method in which go omissions were replaced with the maximum RT (1000ms) (Verbruggen et al., 2019).

#### Computational modelling

To gain insight into whether stimulation affected the processes underlying going and stopping, we fitted the stop accuracy and response time distributions to a dependent process model (DPM) with the Race Against Drift Diffusion toolbox (RADD v0.5.5) (Dunovan et al., 2015; Dunovan & Verstynen, 2019). The DPM assumes that the execution process (*θ_e_* begins to accumulate after a delay (*tr*) until reaching an upper decision threshold (*a*), yielding a go response (Figure 2). The dynamics of *θ_e_* are described by the stochastic differential equation, accumulating with a mean rate of *v_e_* (i.e., execution drift rate) and a standard deviation described by the dynamics of a white noise process (*dW*) with diffusion constant *σ* as follows:

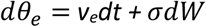

In the event of a stop signal, the braking process (*θ_b_*) is initiated at the current state of *θ_e_* with a negative drift rate (*v_b_*). If *θ_b_* reaches the 0 boundary before *θ_e_* reaches the execution boundary no response is made. The change in *θ_b_* over time is given by:

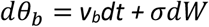

The dependency between *θ_e_* and *θ_b_* in model is implemented by declaring that the intial state of *θ_b_* is equal to the state of *θ_e_*.

To determine if any of the model parameters (execution drift rate (*V_e_*), braking drift rate (*v_b_*), boundary height (*a*) or execution onset delay (*tr*)) changed during stimulation we fitted four models to the average group data, each allowing only one of the parameters to vary for the within subject factors FREQUENCY (20Hz, 70Hz), and STIMULATION CONDITION (ON, OFF). The fitting procedure was aimed at minimizing a cost function equal to the sum of the squared and weighted errors between vectors of observed and simulated response probabilities, stop accuracy and response time quantiles of correct go responses and failed stop responses (error RT). To obtain an estimate of fit reliability for each model, we restarted the fitting procedure from 20 randomly sampled sets of initial parameter values (based on 2000 sampled parameter sets). All fits were initialized from multiple starting values in steps (step size .05) to avoid biasing model selection to unfair advantages in the initial settings. Each initial set was then optimized using a basinhopping algorithm to find the region of global minimum followed by a Nelder-Mead simplex optimization for fine-tuning globally optimized parameter values. The simplex-optimized parameter estimates were then held constant, except for the designated parameter that was submitted to a second simplex run to find the best fitting values for the four conditions. Finally, the model fits were compared in goodness-of-fit with the Akaike information criterion (AIC) and Bayesian information criterion (BIC). A difference of AIC/BIC between 3-10 is considered moderate evidence for one model over the other and >10 as strong evidence (Lee & Wagenmakers, 2014). For more details on model fitting, model code, simulation, cost function weights, and animations see Dunovan et al. 2015, 2019 and https://www.github.com/coaxlab/radd.

### Electrical field modeling

For the last 20 participants, a Philips 3T MRI scanner with a 32-channel head coil was used to acquire high resolution T1 and T2-weighted images, with and without fat suppression (4 scans in total). T1-weighted structural images were acquired using magnetization prepared rapid gradient echo (MPRAGE; TR=9.60ms, TE=4.60ms, 222 sagittal slices, 0.98×0.98×1.2mm voxels). T2-weighted structural images were acquired with TR=2500ms, TE=203ms, 200 sagittal slices, 1.02×1.01×1mm voxels. To simulate the electrical field expansion of the 1-by-4 electrode montage, computational modeling was performed (www.simnibs.org) using a finite element head model derived from the four T1 and T2 scans of each individual (Opitz et al., 2015). All electrodes were modeled as a 2mm thick rubber layer (conductivity 0.1S/m) with a 1mm thick layer of conductive gel underneath (conductivity of 3S/m, as stated by the manufacturer). The positions of the tACS electrodes were determined based on the localization of the EEG electrode positions of all 128 sensors and three landmarks positions (nasion, left and right preauricular). Also the positions of the connectors were explicitly modeled (modeling procedure described in detail in Saturnino et al. (2015)). A current strength of 500μA was simulated, corresponding to 1000μA peak-to-peak amplitude. Finally, the normalized predicted electric field distribution mesh was converted to nifti (https://github.com/ncullen93/mesh2nifti).

To evaluate whether the amount of current that reached the preSMA was related to the effect of stimulation on behavior, the normalized electrical field strength was extracted for the region of interest. This was done as follows: A ROI was created based on peak fMRI activation coordinates (contrast stop>go) in previous studies using the same paradigm (Coxon et al., 2016; Leunissen et al., 2016)(sphere with 10mm diameter centered around the coordinate [11, 10, 62] of the Montreal National Institute (MNI) space). The ROI was transformed into subject space using the inverse of the deformation fields generated by the simnibs pipeline. Average normalized predicted electric field in the resulting individual preSMA ROIs of each session were related to the percent change in force during beta and gamma stimulation using linear regression.

### EEG analysis

EEG data was analyzed using the FieldTrip toolbox for EEG/MEG-analysis (Oostenveld et al., 2011). EEG during no stimulation trials was only analyzed in the last 20 participants due to the high amount of data loss in the first 16 (see tACS-EEG procedures). Pre- and post-tACS resting EEG recordings were available for all 36 participants. Since non-neural signals contaminate the low amplitude gamma-band activity in scalp EEG all analyses only focus on beta-band activity (Muthukumaraswamy, 2013).

#### Pre- and post-tACS resting EEG measurements

Pre- and post-tACS resting EEG recordings were re-referenced to the average reference. Bad channels were rejected upon visual inspection. Subsequently, the data was band-pass filtered at 1-100 Hz. Independent component analysis was used to identify ocular artifacts, which were then projected out of the data. Components were selected based on the highest weights of the mixing matrix contributing to the horizontal and vertical electro-oculogram (see supplementary material). Finally, the data was epoched in 1s segments and bad segments were rejected upon visual inspection. A fast Fourier transformation (FFT) for frequencies between 4 and 45Hz was performed on the first 100 artifact-free segments using a Hanning window and 10s zero-padding.

In order to determine the beta peak frequency, the resulting spectra of the pre-measurement in the 20Hz stimulation session were averaged. The 1/f component was removed by fitting a linear trend (least-squares fit) to the log-transformed spectrum (Haegens et al., 2014; Nikulin & Brismar, 2006). Subsequently, a 3^rd^ order Gaussian curve was fitted to the power spectra to estimate the individual peak frequency.

Changes in absolute beta power from pre-to post-tACS were investigated by calculating the percent change in mean power at 13-30Hz in the averaged pre and post spectra.

#### Task performance in tACS-free intervals

To capture potential differences in event-related synchronization or desynchronization (ERS/ERD) we extracted 2s before and 0.7s after the stop signal presentation. Epochs were re-referenced to the average reference and noisy channels and epochs were rejected upon visual inspection. Independent component analysis was used to identify ocular artifacts, which were then projected out of the data (see resting EEG measurements). EEG data of one participant had to be discarded due to excessive (eye)movements. Subsequently, the data was band-pass filtered between 1-100 Hz. To avoid boundary jumps caused by the filtering procedure the first and last 300ms of the epochs were discarded. Complex Fourier spectra were extracted with Morlet wavelets between 4 and 45 Hz with step size of 0.5 Hz and a fixed width of 7 cycles. The resulting absolute time-frequency spectra were averaged per condition and the conditions were compared with a dependent-sample cluster-based permutation t-test (two-tailed, 5000 permutations, cluster alpha of 0.05). This procedure ensures correction for multiple comparisons over time and frequencies (Maris & Oostenveld, 2007).

Pearson correlations were used to test for relationships between beta activity and behavior (two-tailed, 5000 permutations). The absolute time-frequency spectra were averaged over the full frequency range (13-30Hz). An FDR correction (alpha 0.05) was applied for correcting for testing multiple time points.

### Statistical analysis

All statistical analyses were performed with R version 4.0.0 (R Core Team, 2020) using packages nlme version 3.1-147 (Pinheiro et al., 2020) and multcomp version 1.4-13 (Hothorn et al., 2008).

For the behavioral outcome measures linear mixed effects (LME) models were specified with tACS FREQUENCY (20Hz, 70Hz) and STIMULATION CONDITION (ON, OFF) as fixed factors. For force outcome measures LME models were specified with tACS FREQUENCY, STIMULATION CONDITION (ON, OFF) and TRIAL TYPE (successful stop, go) as fixed factors. For all LMEs random intercepts were modeled on subject level (restricted maximum likelihood criteria, REML). Results for LME models are given as Type III sums of squares for sequentially fitted fixed effects (F, df, p). Significant results from simultaneous pairwise post-hoc comparisons with Tukey contrasts are reported with adjusted *p*-values for estimates of contrasts (estimated mean difference ± SE, *z*-value, adjusted p_Tukey_).

We assessed whether the percent change in force between stimulated and non-stimulated trials was significantly different from zero using a two-tailed one-sample t-test.

A one-tailed Pearson correlation was used to assess the relationship between behavior and normalized electrical field strength.

The percent change scores in beta power from pre-to post-stimulation were non-normally distributed (Shapiro-Wilk test p<0.05). Therefore, a one-sample Wilcoxon Signed Rank Test was used to test whether the percent change in beta power was significantly different from zero. The 20 and 70Hz sessions were compared with the Wilcoxon signed-rank test.

The influence of tACS FREQUENCY (20Hz, 70Hz), SESSION (first, second), and TIME POINT (pre, post) on subjective level of stimulation-related discomfort (VAS_discomfort_) and subjective level of fatigue (VAS_fatigue_) were analyzed with a repeated measures ANOVA.

Descriptive statistics are given as average ± standard deviation unless indicated differently.

## Acknowledgments

This work was supported by the Internal Research Fund KU Leuven (C16/15/070), Research Foundation Flanders (FWO) grants (G089818N, G0F7616N, G093616N, I005018N) and an Excellence of Science grant (EOS 30446199, MEMODYN). IL is supported by an individual fellowship of the FWO (12M6718N) and EU (MSCA 798619). JC is supported by the Australian Research Council (ARC DP200100234 and DP180102066).

