## Supplemental Material for "Effects of beta- and gamma-band rhythmic stimulation on motor inhibition"

**Supplementary Table 1.** Subjective level of discomfort and fatigue based on Visual Analog Scale ratings.

| Subjective rating between 1-10 | | 20Hz session | 70Hz session | Statistics | | | |
|---|---|---|---|---|---|---|---|
| | | | | df, Error df | Main effect of frequency (20Hz, 70Hz) | Main effect of time (pre, post) | Interaction |
| Discomfort | | 1.35±1.8 | 0.66±1.3 | 1,30 | F=2.64 p=0.12 | | |
| Fatigue | pre | 2.40±1.9 | 2.56±1.6 | 1,95 | F=0.01 p=0.93 | **F=48.23 p<0.001** | F=0.46 p=0.50 |
| | post | 4.53±2.1 | 4.34±2.2 | 1,95 | | | |

Mean ± standard deviation is reported.

128-channel electrode map

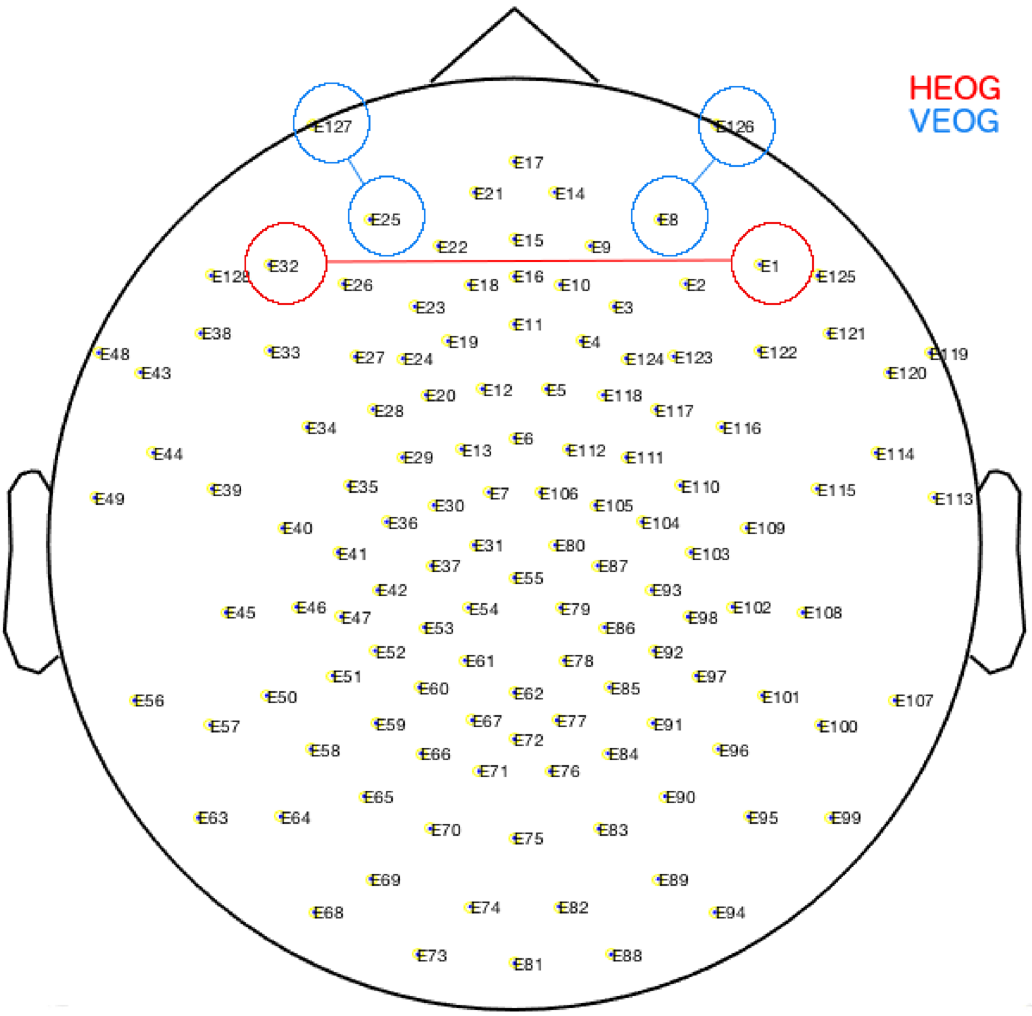


**Supplementary Figure 1.** By linearly combining EEG signals collected from the 128-channel geodesic net we obtained the horizontal electrooculogram (hEOG) and the vertical electrooculogram (vEOG).
